# Characterization of amylase and protease activities in the digestive system of *Concholepas concholepas* (Gastropoda, muricidae)

**DOI:** 10.1101/132100

**Authors:** Alejandro P. Gutierrez, Viviana Cerda-Llanos, Diego Forttes, Nelson Carvajal, Elena A. Uribe

## Abstract

*Concholepas concholepas* (loco) is a carnivorous gastropod that inhabits the coast of Chile and Peru. Its fisheries showed a great importance in the past decades, however, now mainly relies on artisanal management of wild stocks. Feeding is one of the important factors that have restrained the establishment of large scale field rearing of loco. *C. concholepas* food preferences consist of mytilids and cirripeds, however its digestive physiology has not been studied and its digestive enzymes have not been yet characterized. The purification of amylase and protease from the digestive gland and the gland of Leiblein of *C. concholepas* were performed by ionic exchange chromatography (DEAE-cellulose), and substrate-PAGE indicated the presence of the amylase and protease in the fractions collected from the column. Amylase enzymatic assays showed its maximum activity at pH 7.0 and 50°C in the digestive gland. Protease on the other hand showed a great acidic activity, specifically at pH 3.0 in both organs, also at 50°C. Inhibition of the amylolytic activity was observed in the presence of EDTA, and metal ions like MnCl_2_, MgCl_2_, ZnSO_4_, while it was enhanced in the presence of CaCl_2_, NaCl, KCl. Protease inhibition assays were also performed evidencing mainly the presence of aspartic proteases and a low but not inexistent presence of serine proteases. Our results provide evidence of important proteolytic but also amylolytic activities present in the digestive system of loco, providing evidence that this mollusc has wider digestive capabilities than initially though, which could potentially lead to the development of alternative food diets.

## INTRODUCTION

*Concholepas concholepas*, commonly known as loco, is a carnivorous muricid gastropod that inhabits the entire coast of Chile and southern Peru. It was once one of the most important benthic shellfish mollusc extracted in Chile (Castilla and Defeo 2001) but the increasing demand and consequently extraction lead to its overexploitation in the late 80´s (Castilla and Defeo 2001; Leiva and Castilla 2001). Currently its harvest is basically sustained by both artisanal fisheries and the establishment of “Management and Exploitation Areas” (MEAs) which account for approximately 4000 tons per year (SERNAPESCA 2014) and international markets mainly influenced by Asia have maintained the demand for this resource.

The digestive system of loco is similar to that of other gastropods with a well-defined esophagus, intestine and stomach (Huaquín 1966; Maldonado 1965); however, to our knowledge, little information is available with respect to the digestive physiology of this species, although it is known that it is a carnivorous gastropod which food preferences and requirements include Cirripeds, Ascidians and some Mytilids (Stotz et al. 2003, Dye 1991; Méndez and Cancino 1990). To date, breeding of loco has not been achieved successfully due to several factors; among those, feeding has been identified as one of the important issues hindering the establishment of sustainable field rearing (Manríquez et al. 2008).

Nutrition is a costly but important component of the farming process of all species. Due to the predation nature of loco, the problem is substantial because of the need of a coculture of live prey stocks. The formulation of an efficient feeding diet is desirable in most important aquaculture species, therefore, the understanding of their digestive physiology is an essential step in this process (Zambonino Infante and Cahu 2007). Digestive enzymes play an essential role in food degradation and are determinants of the digestibility and assimilation efficiency (Fernández et al. 2001; Picos-Garcia et al. 2000). In fact the ability of an animal to digest and absorb nutrients depends on the presence and the quality of their digestive enzymes (Kumar et al. 2007; Alarcón et al. 1998). In fish the activity of digestive enzymes are thought to indicate the feeding ecology and diet, which in some cases correlates with a plasticity in their production (German et al. 2004; Fernández et al. 2001). In fact they have been a mayor object of study in marine animals because of their biochemicals properties and variables applications (Fu et al. 2005; Chakrabarti et al. 1995; Dimes et al. 1994).

Amylase catalyzes the endohydrolysis of 1,4-α-D-glucosidic linkages of polysaccharides and are widely distributed in animals. Amylases are known to be more active in herbivores than in carnivores, due to a diet composed almost exclusively of carbohydrates (German et al. 2010). In molluscs amylase has been described in herbivorous species such as *Mytilus galloprovincialis* (Lombraña et al. 2005), *Hyriopsis bialatus* (Areekijseree et al. 2004), *Perna viridis* (Sabapathy and Teo 1992) and several Haliotis species (Nikapitiya et al. 2009; Hsieh et al. 2008; Viana et al. 2007; Garcia-Carreno et al. 2003; Tsao et al. 2003; Picos-Garcia et al. 2000). Further, it has also been described in carnivorous fishes of great aquaculture importance like salmonids, but showing lower levels of amylase activity compared to proteolytic activity (Hidalgo et al. 1999). Digestive proteases such as trypsin, chymotrypsin carboxypeptidases and aminopeptidases, are mainly produced in the digestive gland, but also in the gland of Leiblen which is highly developed in muricids gastropods (Andrews and Thorogood 2005; Mansour-Bek 1934). In fact, the Gland of Leiblein has been described as a secretor of proteolytic enzymes and a useful defense barrier (Ponce et al. 1990) in *C. concholepas*. Proteolytic enzymes play a fundamental role in food digestion and storage of chemical energy (Sainz et al. 2004), considering that they are in charge of the degradation of proteins obtained from the diet. Moreover, since protein utilization is fundamental for growth, proteases have an important role to play in fish development (Kumar et al. 2007). Proteolytic enzymes in the digestive organs of invertebrates have been well documented and characterized, e.g. spinny lobster *Panulirus interruptus* (Celis-Guerrero et al. 2004), shrimps like *Artemesia longinaris* (Fernández Gimenez et al. 2002), *Penaeus indicus* (Omondi and Stark 2001), the mud crab *Scylla serrata* (Pavasovic et al. 2004), the pacific oyster *Crassostrea gigas* (Luna-Gonzalez et al. 2004) and various species of abalone as red abalone *Haliotis rufescens* (Garcia-Esquivel and Felbeck 2006), green abalone *Haliotis fulgens* (García-Carreño et al. 2003), black abalone *Haliotis rubra* (Edwards and Condon 2001) and in juvenile South African abalone *Haliotis midae* (Knauer et al. 1996), however, studies of the digestive enzymes in carnivorous molluscs are scarce.

The aim of this study was to analyze for the first time the amylase and proteolytic activities present in the digestive system of the carnivorous gastropod *C. concholepas*. We aimed to evaluate the potential ability of the animal to assimilate and digest protein and carbohydrates regardless of its normal food preferences. These analyses could provide significant information for the understanding of their carnivorous nature and ability to digest food, but also provide insights on the possible formulation of new feeding diets.

## MATERIALS AND METHODS

### Sampling of C. concholepas

The *C. concholepas* individuals were collected by fishermen from a legally authorized extraction zone in the shores of Maule fishing cove, located in the eighth region of Chile (-37.009,-73.190). The individuals were chosen with respect to size as they were collected by the fishermen.

### Preparation of extracts and partial purification of enzymes

Individuals were dissected and their digestive system were extracted, specifically the digestive gland and the gland of Leiblein. Once isolated, tissues were washed with distilled water, weighed and then homogenized in 10 mM Tris-HCl buffer, pH 7.5 at 4°C. To eliminate feed residues, solid material and lipids, the homogenate was centrifuged at 10.500 X *g* for 10 min, also at 4°C. The supernanant crude extract was dialyzed for at least 3 hours at 4°C in 10mM Tris-HCl, pH 7.5 buffer, then the extract was applied trough a anionic exchange chromatography of diethylaminoethyl-cellulose (DEAE-cellulose) calibrated with Tris-HCl 10 mM, pH 7.5. 50 ml of the crude extract was applied into the column several times in order to allow the proteins electrostatically interact with the resin. The column was washed with 10 mM Tris HCl buffer, pH 7.5 and then eluted in two steps: first with 250 mM KCl buffer in 10 mM Tris-HCl, pH 7.5, followed by 500 mM KCl buffer in 10 mm Tris-HCl, pH 7.5. Fractions of approximately 1 ml were collected, and then used for further analysis.

### Amylase Enzyme assays

Amylase activity was determined according to Bernfeld (1955). Briefly, 50 μl of the enzyme was mixed with 50 μl of 100 mM Tris-HCl buffer pH 7.0 and 100 μl of 1% starch at pH 7.0. The mixture was incubated at 40°C for 30 minutes, followed by the addition of 200 μl dinitrosalicylic acid (DNS) to stop the reaction. After mixing by swirling, the mixture was heated for 5 minutes at 100°C to allow the DNS reaction. Afterwards, 2 ml of distilled water were added, and the absorbance was measured at 540 nm. One unit of activity was defined as the amount of maltose released in 1 minute of enzymatic reaction.

For the determination of optimum pH of amylase, the assay was conducted using the following buffers with variable pH values: 0.1M KCl pH 2.0; 0.1M citrate pH 3.0, 4.0 and 5.0; 0,1M phosphate pH 6.0, 7.0 and 8.0; 0.1M Gly-NaOH pH 9.0, 10.0 and 11.0. The determination of amylase activity was carried out as described earlier, plus the addition of 300μl of the selected buffer to the reaction mixture. The temperature effect on amylase activity was determined by assaying enzyme activities at temperatures of incubation ranging from 10 to 70°C, at optimum pH.

To determine the effect of metal ions in amylase activity, the protein extract was first dialyzed in 5 mM Tris-HCl, pH 7.5 for 2 hours and then the amylase activity was measured with: 10 mM of CaCl_2_, NaCl, KCl, ZnSO_4_, MnCl_2_, MgCl_2_, and the chelating agent ethylenediaminetetraacetic acid (EDTA).The assay was carried out at 40°C, pH 7.0.

### Proteolytic Enzymes assays

Proteolytic activity was determined by the digestion of casein according to (Glass et al. 1989), with modifications. The assay was conducted using the following buffers with variable pH values: 0.1M KCl (pH 2.0), 0.1M citrate (pH 3.0 - 5.0), 0.1M phosphate (pH 6.0 - 8.0), 0.1M Gly-NaOH (pH 9.0 - 11.0). Briefly, 50 μl of the enzyme was mixed with 200 μl of the buffer according to pH and 50 μl of 2% casein at pH 7.0. The mixture was incubated for 30 minutes at 37°C. The reaction was stopped by adding 250μl of 10 % (w/v) trichloroacetic acid (TCA) and left at 4°C for 30 min. The samples were then centrifuged at 10.000 rpm for 5 minutes and 350 μl of the supernatant was mixed with 900 μl of 0.5 M sodium carbonate (Na_2_CO_3_) followed by the addition of the Folin-Phenol reagent. The absorbance of the reaction mixture was recorded at 660 nm to quantify the amount of tyrosine produced measured in units (U).One U is defined as the amount of tyrosine released in 30 minutes of enzymatic reaction. The specific activity of both enzymes is defined as units per minute per mg protein (U min^-1^ mg prot.^-1^).

In order to determine the presence of major classes of proteolytic enzymes, the effect of inhibitors on the proteolytic activity was tested according to García-Carreño (1992). Enzyme extracts were incubated with different specific proteinase inhibitors. Phenylmethylsulfonyl fluoride (PMSF, 100 mM in ethanol) were used as inhibitors of proteinases belonging to the serine class. Ethylenediaminetetraacetic acid (EDTA, 500mM in distilled water) was used as an inactivator of metallo-proteinases. For inhibition of aspartic proteases, 5μg/ml of Pepstatin A (50 μg /ml in DMSO or methanol) were used. Proteolytic activity was measured as described above.

### Electrophoresis

The fractions collected from DEAE-column with amylase and protease activity were analyzed in 12% SDS-PAGE according to Laemmli (1970). The samples were mixed with a solution of electrophoresis buffer which contained 1M Tris HCl buffer, 10% SDS, 10% glycerol, 0.5% bromophenol blue and 10% β-mercaptoetanol in a 1:5 proportion. After boiling the samples for 5 minutes, the electrophoresis was carried out at 80 V for at least two hours. The gels were stained in 0.1% Comassie brilliant blue R-250.

Substrate-SDS-PAGE for detected amylase activity was performed on 4% stacking gel and a 12% resolving gel in presence of 2% starch for detection of amylase activity. Electrophoresis was carried out at 60 V until the samples passed through the stacking gel, and then raised to 90V for at least three hours. The samples and the electrophoresis buffer were mixed in a 1:1 v/v proportion, in absence of reducing agents. The amylase activity was detected by staining the gels with iodine after incubation in 100 mM Tris-HCl pH 7.5 for 2 h at 40°C.

Protease Substrate-PAGE was performed on 4% stacking gel and an 8% resolving gel copolymerized with 2% casein for detection of proteolytic activity. Electrophoresis was carried out at 70 V until the samples pass through the stacking gel, and then raised up to 100V for at least three hours. The samples and the electrophoresis buffer were mixed in a 1:1 v/v proportion, in absence of reducing agents and SDS and without boiling. The proteolytic activity was detected by staining the gels with 0.1% Commassie brilliant blue G-250 after incubation in 50mM Tris HCl pH 7.0 and 0.1M Citrate buffer pH 3.0 at 37°C overnight.

### Protein content

Protein content was determined by the method of Bradford (Bradford 1976) using bovine serum albumin as standard.

## RESULTS

### Enzyme purification

The amylase and protease enzymes were purified through of a DEAE-cellulose column calibrated to pH 7.5. The enzymes were retained in the column and then eluted with 250 mM and 500 mM KCl in 10 mM Tris-HCl, pH7.5. The profile of total proteins and amylase activity are shown in Table 1.

**Table 1.**
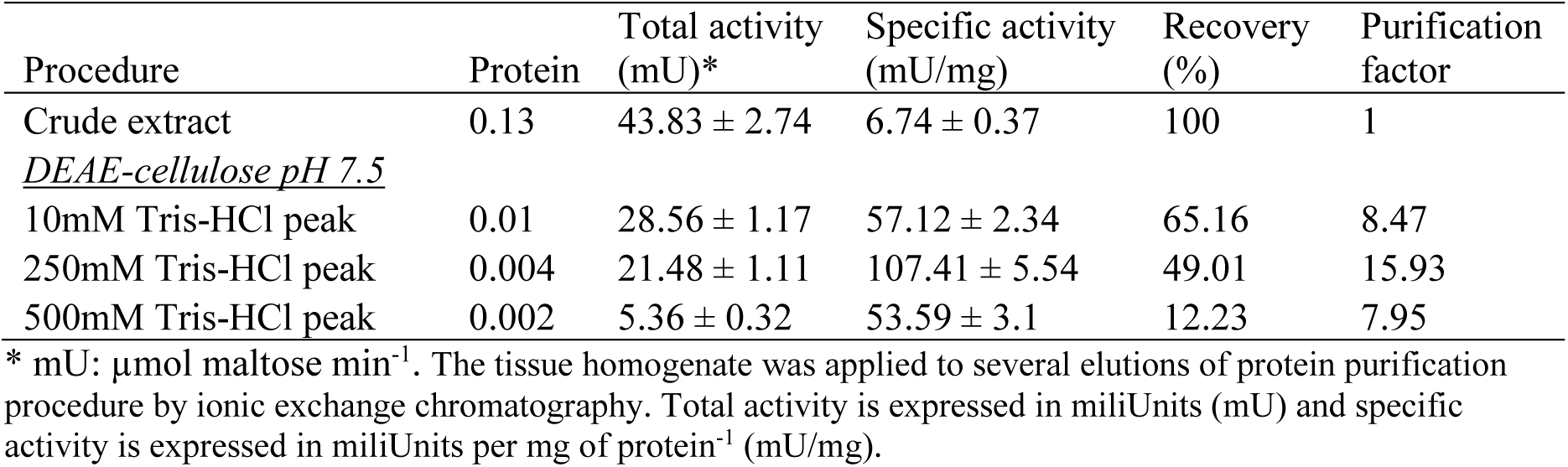
Summary of the purification procedures and enzymatic activity of amylase from *C. concholepas*

The amylase activity was also expressed in specific activity (mU/mg) that increased according to its purification. The crude extract from the digestive gland showed a specific activity of 6.74±0.37 mU/mg. After the extract was applied to the DEAE-cellulose column purification, the specific activity of amylase increased up to 57.12±2.34 mU/mg in the 10mM Tris HCl buffer wash. The following elutions with 250 and 500mM KCl buffer, showed a specific activity of 107.41±5.54 mU/mg and 53.59±3.1 mU/mg respectively. The gland of Leiblein did not show any amylase activity.

On the other hand, the digestive gland showed slightly higher proteolytic specific activity (7.90±0.31 U/mg prot^-1^) than the gland of Leiblein (6.81±0.37 U/mg prot^-1^) in the crude extract (Table 2), however, it was much higher in the last step of the purification procedure showing an increment of approximately 60% in relation to the 13% of difference between them in the crude extract.

**Table 2.** Total and specific protease activity in the gland of Leiblein and the digestive gland of C*. concholepas*.

| Procedure | Digestive Gland |  | Gland of Leiblein |  |
| --- | --- | --- | --- | --- |
|  | Total activity (mU)* | Specific activity (mU/mg) | Total activity (mU)* | Specific activity (mU/mg) |
| Crude extract | $67.11 \pm 2.63$ | $7.90 \pm 0.31$ | $44.27 \pm 2.42$ | $6.81 \pm 0.37$ |
| <u>DEAE-cellulose pH 7.5</u> |  |  |  |  |
| 10mM Tris-HCl peak | $49.37 \pm 3.07$ | $32.92 \pm 2.04$ | $34.45 \pm 1.93$ | $28.71 \pm 1.72$ |
| 250mM Tris-HCl peak | $35.30 \pm 2.12$ | $117.66 \pm 7.06$ | $19.36 \pm 1.12$ | $75.94 \pm 4.39$ |
| 500mM Tris-HCl peak | $30.57 \pm 1.94$ | $203.81 \pm 12.9$ | $17.29 \pm 1.56$ | $123.52 \pm 11.14$ |

\* mU: $\mu\text{mol tyrosine min}^{-1}$ . The tissue homogenate was applied to several elutions of protein purification procedure by ionic exchange chromatography. Total activity is expressed in miliUnits (mU) and specific activity is expressed in miliUnits per mg of protein<sup>-1</sup> (mU/mg).

Specific activity was low in the crude extract, however it consistently increasing from 7.90±0.31 U/mg prot^-1^ to 203.81±12.9 U/mg prot^-1^ in the digestive gland and from 6.81±0.37 U/mg prot^-1^ to 123.52±11.14 U/mg prot^-1^ in the gland of Leiblein during the purification procedure.

### Sustrate SDS-PAGE

Substrate SDS-PAGE results (Figure 1a), shows the presence of amylase activity on two bands in selected fractions from the 250mM KCl buffer elution, detected by means of degradation of starch contained in the gel. This could represent the presence of two possible isoforms of purified amylase. These results are in concordance with SDS-PAGE analysis (Figure 1b), where the same fractions shows the presence of two bands, similar as the ones seen in Figure 1a.

**Figure 1.**
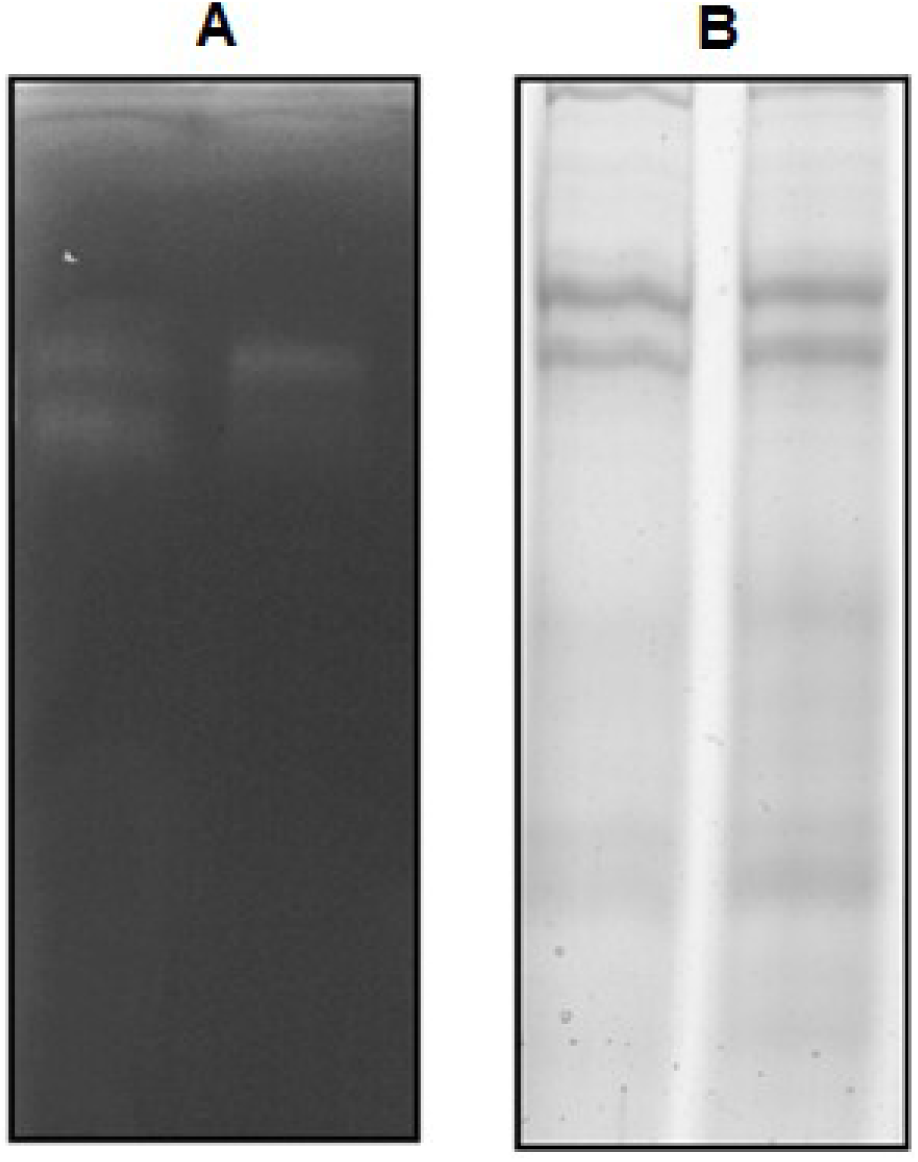
Electrophoretic analysis of two purified fractions of 250mM KCl buffer elution from DEAE column. **A.** Substrate SDS-PAGE. The presence of two bands in every fraction shows amylase activity and two possible isoforms of the enzyme. **A.** SDS-PAGE. The presence of two main bands in the same fractions corroborate the presence of amylase and the two isoforms like shown in A).

In the case of proteases, substrate PAGE revealed the presence of white bands in a dark background which indicate the existence of proteolytic activity in fractions from the DEAE column in the gland of Leiblein and the digestive gland (Figure 2). This technique showed clear bands when the gels were incubated at pH 3.0 which is in agreement with the results obtained earlier, where the highest proteolytic activity was found at this pH. In contrast, at pH 7.0 the bands were almost null but not inexistent. Proteolytic activity was found at this pH, but in a much lower level than the acidic pH.

**Figure 2.**
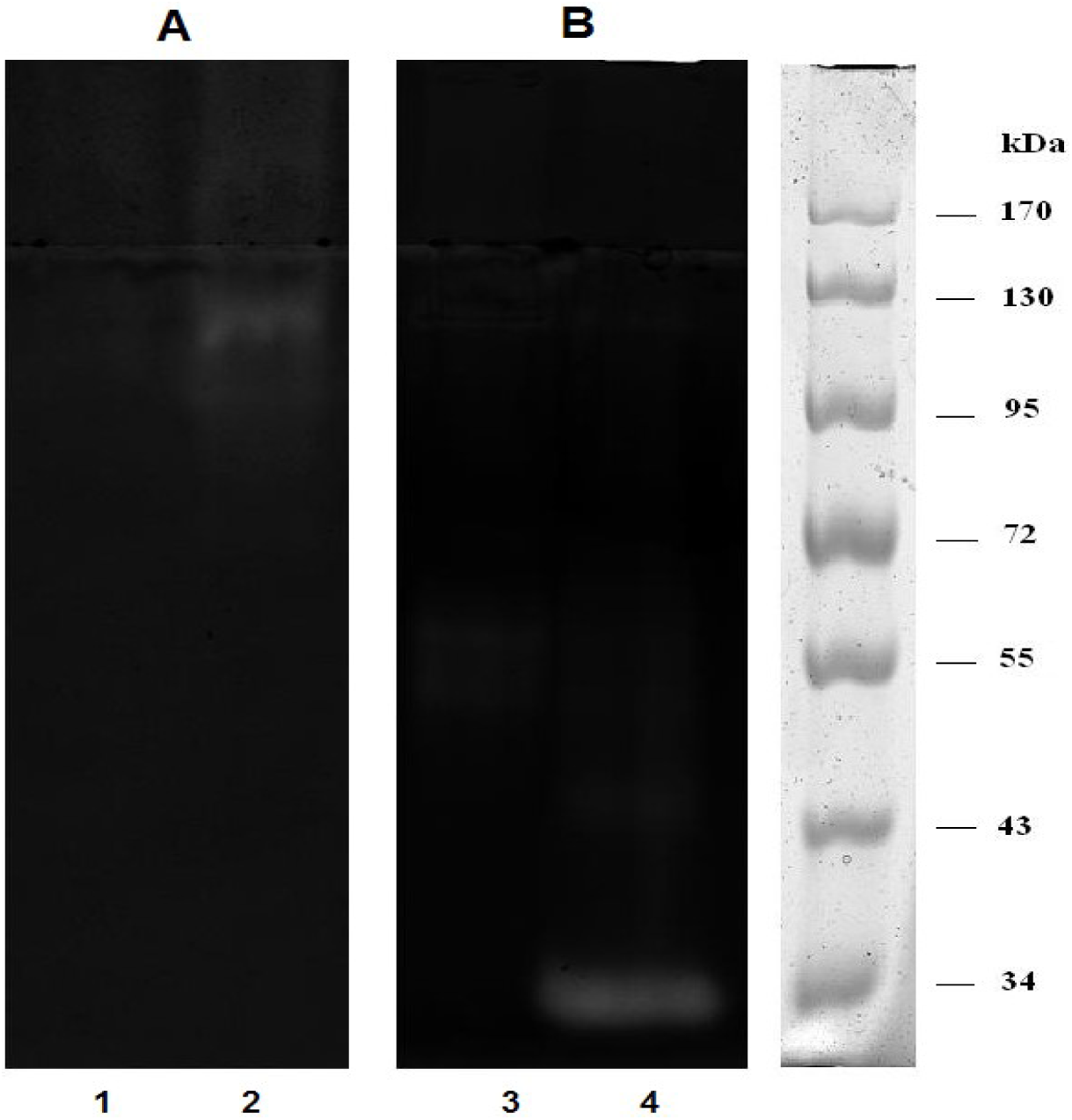
Substrate-PAGE shows proteolytic activities in fractions from the 250mM KCl elution of the DEAE column at a) pH 7.0 and b) pH 3.0. Lane 1, 3: Digestive gland. Lane 2, 4: Gland of Leiblein of *C.concholepas*.

### Effect of temperature and pH

The effect of pH on the amylase activity was examined at 40°C using different types of buffers for each pH. Analyses showed that the optimum pH for amylase activity was pH 7.0 in the digestive gland (Figure 3). On the other hand, the amylase activity at acidic and alkaline pH was weak but comparable between each other, except for pH 11-12 where the activity dropped drastically, although acidic forms of amylase seem to be more active than alkaline forms.

**Figure 3.**
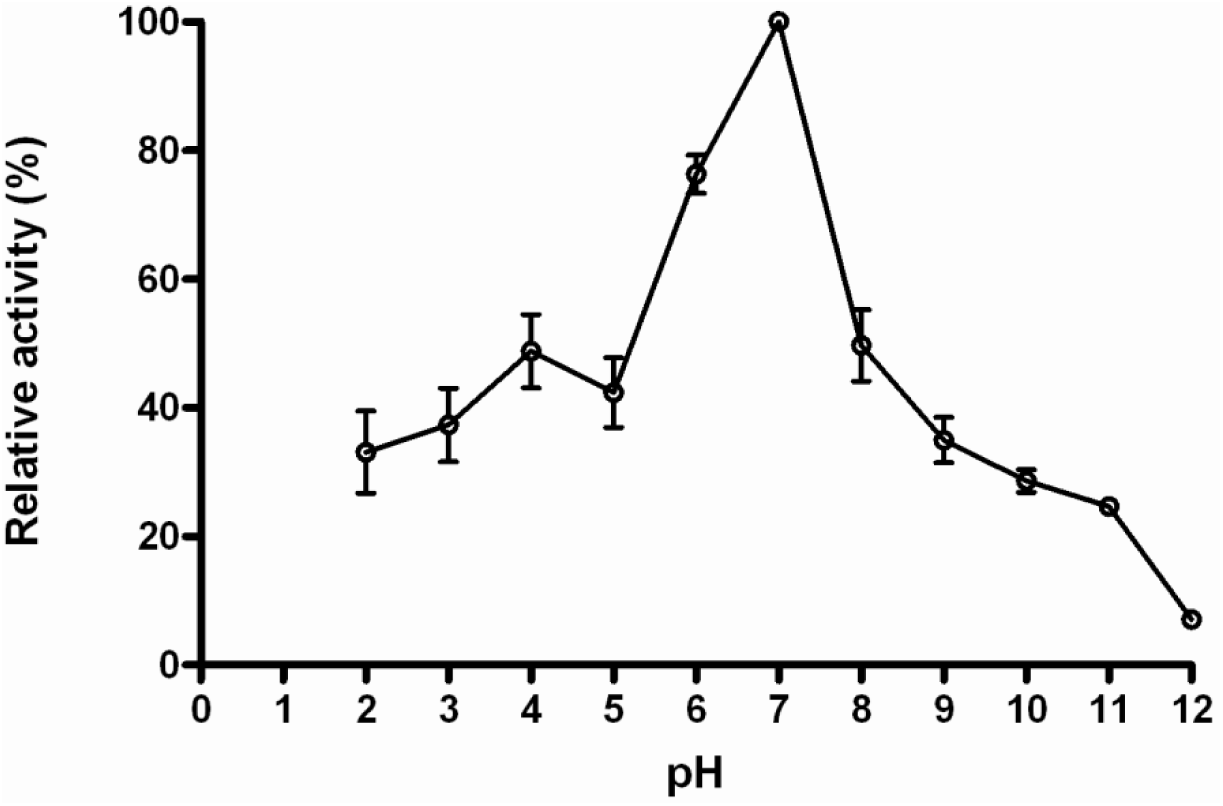
Effect of pH on amylase activity of *C. concholepas*. Maximum activity was detected at pH 7.0 at 40°C for 30 minutes. The assay was repeated 3 times. Error bars represent standard error (±SE).

The highest proteolytic activity in the digestive gland of *C. concholepas* was found at pH 3.0, suggesting that this gland possess a high level of acidic proteolytic enzymes. (Figure 4). At pH 4.0, the activity dropped considerably and was maintained throughout the neutral pH to then slightly rise again in presence of an alkaline medium, although it is not comparable with the high levels of activity at acidic pH. In addition, the gland of Leiblein had an optimum pH of protease activity very similar to the digestive gland showing high activities at pH 3.0, however at neutral pH (6.0 and 7.0) the activity was slightly higher than in the digestive gland. At an alkaline pH, the proteolytic activity was low.

**Figure 4.**
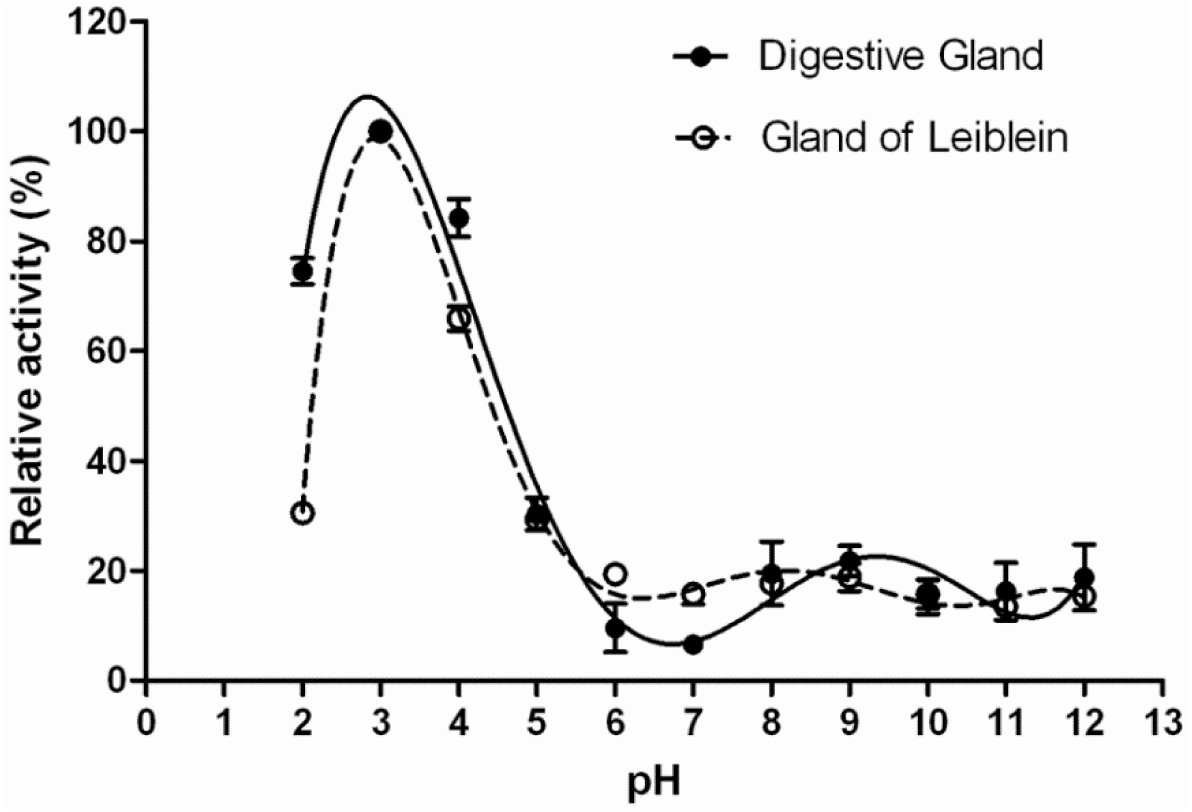
Effect of pH on the proteolytic activity in the digestive gland and the Leiblein gland of *C. concholepas* assayed at 37°C for 30 minutes. Maximum proteolytic activity was found at pH 3.0 in both organs. Error bars represent standard error (±SE).

Amylase activity was measured at different temperatures at pH 7.0. The maximum activity was found at a temperature of 50°C as shown in Figure 5. The activity was retained up to more than 50% of the maximum between 30–60 °C, and then it began to decrease maintaining only 20% of the maximum activity at extreme temperatures.

**Figure 5.**
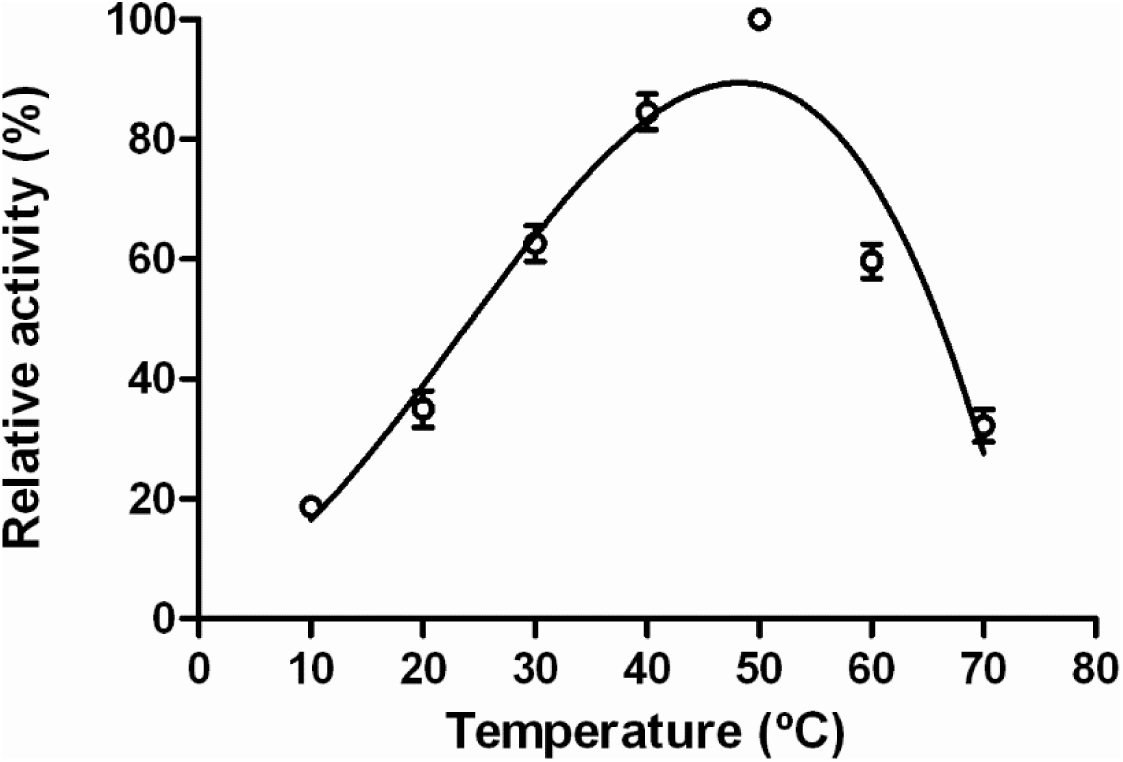
Effect of temperature in amylase activity of *C. concholepas* at pH 7.0. Maximum activity was detected at 50°C in a 30 min. reaction. The assay was repeated 3 times. Error bars represent standard error (±SE).

The effect of temperature on proteolytic activity was also analyzed (Figure 6). The highest activity was detected at 50°C pH 3.0 in both digestive gland and gland of Leiblein, however, at 70°C the enzymes did not inactivate, showing a 9.28% and 20.03% more proteolytic activity than the values detected at 10°C in the gland of Leiblein and the digestive gland respectively.

**Figure 6.**
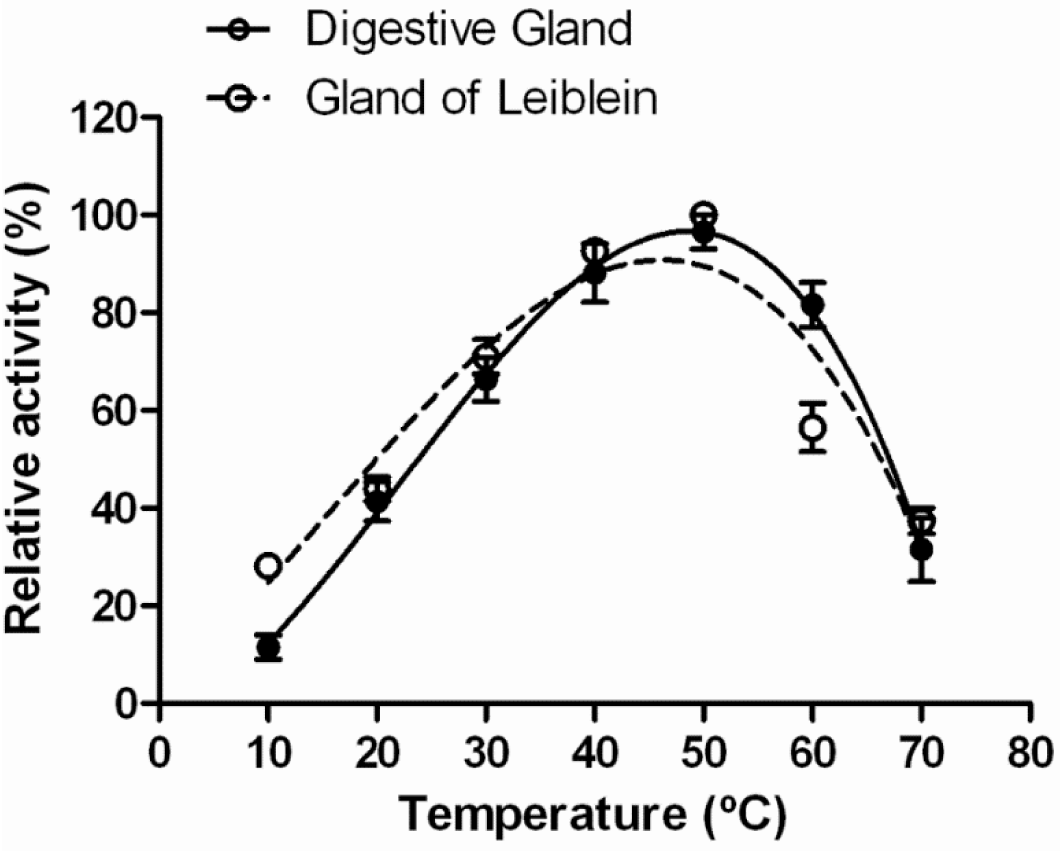
Effect of temperature on proteolytic activity in the digestive gland and in the gland of Leiblein of *C. concholepas* assayed at 37°C for 40 minutes at pH 3.0. Maximum proteolytic activity was found at 50°C in both organs. Error bars represent standard error (±SE).

### Effect of metal ions and inhibitors on amylase and proteolytic activity

Figure 7 shows the effect of metal ions (10mM) on the activity of amylase. The results obtained in the analyses performed in the digestive gland showed an increase on the amylase activity when the enzyme interacts with NaCl, KCl and CaCl_2_, while in presence of EDTA, MnCl_2_, MgCl_2_ and, ZnSO_4_ the activity showed a decrease in the activity.

**Figure 7.**
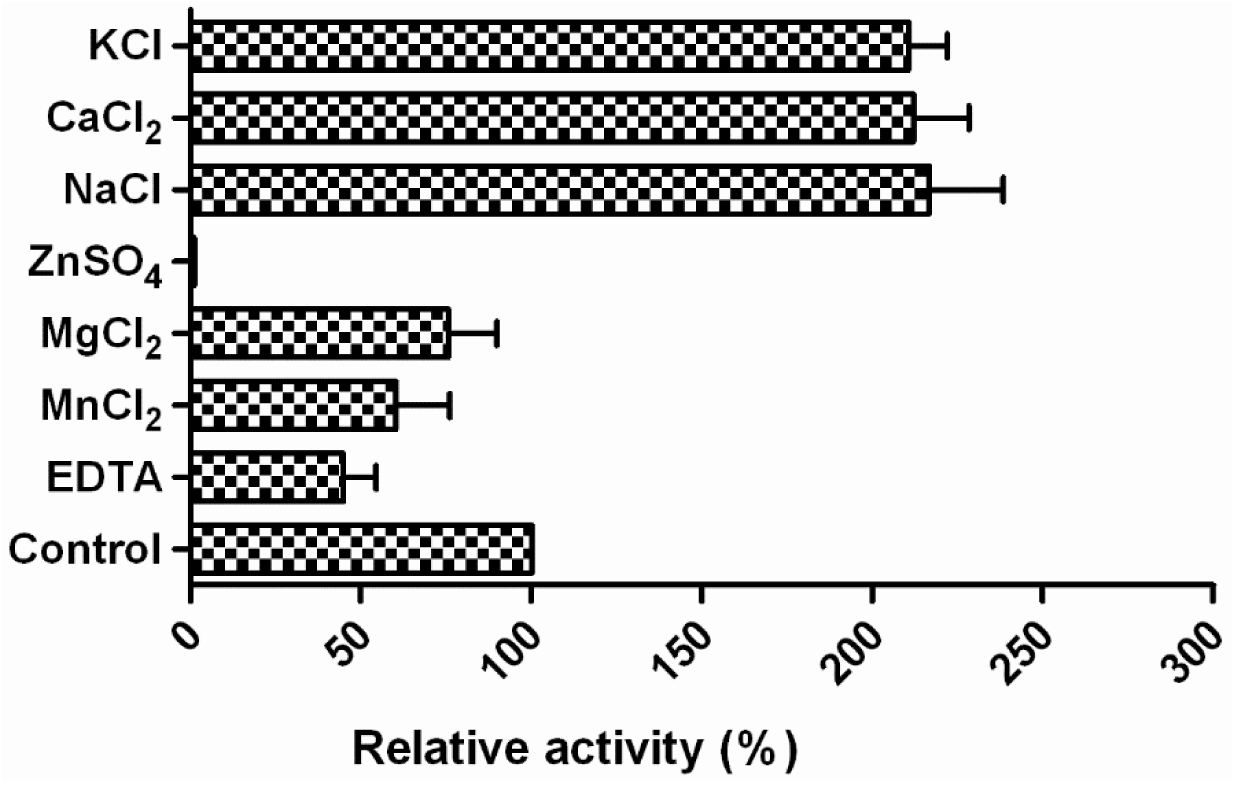
Effect of 10mM of metal ions on amylase activity of *C. concholepas*, at pH 7.0 for 40 min. Activity was enhanced in the presence of NaCl,CaCl_2_ and KCl, and inhibited by EDTA, MnCl_2_, MgCl_2_ and ZnSO_4_. The error bars represent ±SE.

In presence of Na^+^, Ca^2+^ and K^+^, the amylase activity rose over two hundred percent in relation to the control, while when the enzyme was incubated with EDTA and metals ions like Mn^2+^, Mg^2+^ the activity decreased between 50 and 70% approximately. Zn^2+^ greatly affected the amylase activity, producing a decrease in the activity of almost 100% on its presence. The control represents 100% of the total amylolytic activity in absence of metal ions.

Specific protease inhibitors were assessed in the digestive gland and the gland of Leiblein of *C. concholepas*. Table 3 shows the inhibition percentage of the proteolytic activity on these organs. Results show that Pepstatin A inhibited protease activity 100% in the digestive gland whereas in the gland of Leiblein it showed a 90% of inhibition.

**Table 3.** Effect of specific inhibitors on protease activity in the digestive gland and the gland of Leiblein of *C. concholepas.*

| Inhibitor | Final Concentration (mM) | Inhibition (%) |  |
| --- | --- | --- | --- |
|  |  | Digestive Gland | Gland of Leiblein |
| Control | 0 | 0 | 0 |
| Pepstatin A | 5 | 100 | 90.49 |
| PMSF | 5 | 34.4 | 58.62 |
| EDTA | 5 | 33 | 20.77 |

These results provide evidence of the presence of aspartic proteases in both tissues, being pepsin the most probable since Pepstatin A is specific for this enzyme. PMSF was more efficient in the gland of Leiblein inhibiting the activity more than 50%, which might indicate the presence of serine proteases. EDTA inhibited 75% of the proteolytic activity in the digestive gland, whereas in the gland of Leiblein, it only decreased it only in a 40 percent. This also provides evidence of the presence of metalloproteases in the both organs; however, the digestive gland seemed to have the more of these types of proteases due to its high level of inhibition in comparison with the gland of Leiblein.

## DISCUSSION

This study is the first to characterize both protease and amylase activities present in the digestive system of a carnivorous gastropod as *C. concholepas.* Moreover, this study provides novel evidence of the presence of amylase in its digestive system, suggesting wider digestive capabilities for the digestion of food.

### Amylase activity

The purification and characterization of the amylase activities in the digestive gland of *C. concholepas* was carried out by anionic exchange chromatography (DEAE-cellulose calibrate pH 7.5), which was an efficient methodology for the *C. concholepas* amylase purification. Based on its chromatographic behavior, it is possible to suggest that the isoelectric point of the amylase is over 7,5 pH, and this information is relevant for future purifications.

The *C. concholepas* amylase showed an increase on its specific activity during the purification procedure raising up to 107.41±5.54 mU/mg on the 250mM KCl Tris-HCl elution, however its values are lower compared to other gastropod species that are known to be herbivorous and showed higher amylase specific activity (Lombraña et al. 2005; Tsao et al. 2003, 2004). Nevertheless, as a carnivore, lower levels of amylase activity could be expected when compared to the proteolytic activity as it has been described in fishes with different feeding behaviors, but mainly in carnivores (Matus de la Parra et al. 2007; Fernández et al. 2001; Hidalgo et al. 1999; Chakrabarti et al. 1995; Sabapathy and Teo 1993).

The electrophoretic analysis carried out to detect the purified amylase through substrate SDS-PAGE and SDS-PAGE revealed the presence of two bands with both techniques used. These two bands could indicate the presence of two isoforms of the enzyme in the digestive gland of *C. concholepas*. Same number of isoforms have been described for other species where this type of techniques were useful to detect the presence of the enzyme (Fernández et al. 2001). Moreover, these results are in line with other amylase isoforms previously identified in other molluscs species like *Mytilus galloprovincialis* (Lombraña et al. 2005), and small abalone *Sulculus diversicolor aquatilis* (Tsao et al. 2003), and fish like *Pagrus pagrus, Pagellus erytrhinus, Pagellus bogaraveo, Boops boops* and *Diplodus annularis* (Fernández et al. 2001).

The analysis on the effect of temperature on amylase activity at pH 7.0, indicated a maximum activity at 50 °C. These findings agree with previous studies of amylase in abalone (Nikapitiya et al. 2009; Hsieh et al. 2008; Tsao et al. 2003). Our results also concur with the amylase activities reported in other molluscs with optimal temperatures between 30 to 60°C (Albentosa and Moyano 2009; Lombraña et al. 2005; Areekijseree et al. 2004; Tsao et al. 2004; Sabapathy and Teo 1992). The physiological conditions for amylase activity in loco occur around 18 °C and at this temperature the amylase conserved 50% of its maximum activity. In accordance to our findings, an increase of physiological processes is commonly seen at high temperatures in marine invertebrates (González et al. 1990). The amylase from *C. concholepas* showed evidence of a high thermostability within all the range of analyzed temperatures without inactivation detected, being this results in contrast to what was found in other molluscs that showed amylase inactivation after heating over 50°C (Tsao et al. 2003, 2004). Although the *C. concholepas* amylase showed an optimum activity of 50°C, it also showed a remarkable stability at lower temperatures including 10-20°C which are the common temperatures of the waters that the organism inhabits.

In the case of pH conditions, our results concur with those previously described in other molluscs species like abalone (Nikapitiya et al., 2009; Hsieh et al., 2008), mussels (Lombraña et al. 2005; Areekijseree et al. 2004; Fernández-Reiriz et al. 2001) and clams (Albentosa and Moyano 2009). In most studies, amylases have been shown to display a high sensitivity to low and high pH, which is indicative of the need of neutral pH conditions for the digestion of carbohydrates, as it was suggested by Sabapathy and Teo (1992). This also in agreement with the near neutral pH values measured in the digestive systems of some fishes and molluscs (Munilla-Morán and Saborido-Rey 1996; Sabapathy and Teo 1992). Comparative studies are difficult to perform due to the carnivorous nature of loco among molluscs, but in carnivorous fishes amylase activities have been reported (Fernández et al. 2001; Hidalgo et al. 1999; Munilla-Morán and Saborido-Rey 1996). Although the role of amylase in carnivorous species still remains unclear due to the lack of major carbohydrate ingestion (Natalia et al. 2004), the physiological roles of amylases have been hypothesized in relation to their possible function in glycogen utilization as energy source (Nikapitiya et al. 2009; Koyama et al. 2001). The amylase activity in fish depends on the natural diets of each species, having omnivorous and herbivorous species more amylase activity than carnivores (German et al. 2004; Fernández et al. 2001; Hidalgo et al. 1999; Ugolev et al. 1983; Kapoor et al. 1976), which as expected is associated to higher carbohydrate consumption and absorption.

Besides physical factors like pH and temperature, chemical factors, like metal ions have been reported to enhance or inhibit amylase activity (Lombraña et al. 2005; Baker 1983; Trainer and Tillinghast 1982). NaCl for instance, is a monovalent ion that increases amylase activity (Harris et al. 1986), and its effect has been widely described in molluscs (Nikapitiya et al. 2009; Hsieh et al. 2008; Lombraña et al. 2005; Tsao et al. 2004). Our results are in accordance with previous literature, showing an increase on its activity when exposed to 10mM NaCl.. Optimum concentrations of NaCl ranging from 0.01 to 1M, have been described for other marine species (Lombraña et al. 2005; Trainer and Tillinghast 1982; Robson 1979; Wojtowicz and Brockerhoff 1972)

The effects of CaCl_2_ on amylase activity were consistent and similar to the effects of Na^2+^. In fact, a group of amylases have been described as metalloproteins that require Ca^2+^ for their activity and stability levels (Fischer 1960) and its removal may produce a decrease in activity and a lack of thermostability (Violet and Meunier 1989). In addition, Lombraña et. al (2005) described an optimum amylase activity occurring at CaCl_2_ concentrations of 15 and 40 mM in *Mytilus galloprovincialis*. Moreover, in *Haliotis sieboldii* the maximum activity was registered at 10 mM CaCl_2_ (Hsieh et al. 2008). On the other hand, the addition of Mn^2+^ had a negative effect on amylase activity, with a 50% decrease of activity compared to controls. It is known that Ca^2+^ and Mn^2+^ ions could stabilize and activate the enzyme, but can also inhibit it if present at high concentrations (Witt and Sauter 1996).

In the presence of Zn^2+^, the activity of amylase was completely inhibited as it showed a 100% decrease, similar to what have been reported in other species. This inhibition could be due to the direct actions of Zn^2+^ upon the active center of the enzymes (Golovanova 2010). In juvenile perch and roaches intestines, the inhibition of digestive carbohydrases in the presence of Zn^2+^ decreases the rates of carbohydrates assimilation, which in turn influences the efficiency of fish feeding in a negative way (Golovanova 2010). EDTA is a chelating agent and a well-known protease inhibitor, specifically for metalloproteases (Córdova-Murueta et al. 2003), but in this study, it also played a role as an amylase inhibitor, since the activity of amylase decreased in the presence of the agent, which removes calcium and other metal ions from solutions.

### Protease activity

The digestive gland showed slightly higher specific proteolytic activity (7.90±0.31 U/mg prot^-1^) than the gland of Leiblein (6.81±0.37 U/mg prot^-1^) in the crude extract, however, it was much higher in the last step of the purification procedure showing an increment of approximately 60% in relation to the 13% of difference between them in the crude extract. This was expected because this kind of chromatography lowers down the presence of proteins in the fractions collected in every elution, making the activity more specific and higher since there is a less amount of proteins. These results could also indicate that the digestive gland is the main source of proteolytic enzymes as described in marine gastropods from the genus Buccinum where the main source of gastric enzymes is the digestive gland. The gland of Leiblein facilitates digestion through the intestine and the stomach, which is responsible of providing the optimal conditions for proteolytic activity at acidic pH. (Andrews and Thorogood 2005).

The digestive tract of *C. concholepas* showed a high acidic proteolytic activity when exposed to variable pH medium, reporting the maximum proteolytic activity at pH 3.0 in the digestive gland and the gland of Leiblein, which may indicate the presence of aspartic proteases in both organs. Previous studies have described the presence of these kind of proteases in molluscs like green abalone and fishes like tilapia at pH 2.0 (García-Carreño et al. 2003; Yamada et al. 1993) and at pH 3.0 in spiny lobster (Celis-Guerrero et al. 2004). Proteolytic activity was measured at different temperatures showing higher actvity at 50°C in the digestive gland and the gland of Leiblein, just in the range of temperatures that has been described in two sparid fishes gilthead seabream (*Sparus aurata*) and common dentex (*Dentex dentex*) where maximum acidic activity was found between 45-55°C (Alarcón et al. 1998) and in pearl mussel *Hyriopsis bialatus* where was found at 40°C (Areekijseree et al. 2004) under the same pH conditions. This result may suggest that the *C. concholepas* proteases works at suboptimal temperature conditions, due to the high activity found at 50°C compared to the physiological temperature which is around 18°C. This could be explained by arguing a better suited structure for activity at high temperatures which promotes a better enzyme-substrate interaction (Diaz-Tenorio et al. 2006; Whitaker 1994).

Further characterization of the crude enzyme was achieved with specific protease inhibitors. Proteases in the digestive tract of *C. concholepas* seem to belong to two main proteinase classes; aspartic proteinases and serine proteinases (EC 3.4.21.x). This is explained, in the case of aspartic enzymes, by the total inhibition of proteolytic activity in the digestive gland and for the most part in the gland of Leiblein, in presence of Pepstatin A; a very specific inhibitor, with one of the lowest known Ki for pepsin (Castillo-Yañez et al. 2004). The level of inhibition of the acidic proteolytic activity in the gut of *C. concholepas* by Pepstatin A suggests the presence of the aspartic protease pepsin.

For the PMSF assay, the activity of proteases was inhibited in a 34 % in the digestive gland and in a 58 % in the gland of Leiblein of *C. concholepas*, which concur with other similar studies using PMSF on other aquatic species (Dimes et al. 1994; Garcia-Carreño and Haard 1993). This finding indicated that the slight proteolytic activity in the gut of this mollusc at pH 7.0 is due to serine proteases, especially in the gland of Leiblein. In the presence of EDTA, proteases were slightly inhibited which might indicate the existence of some metalloproteases. The low inhibition effect of this chelator on proteases may be explained by the fact that the acidic protease pepsin is typically characterized by a cation-independent mechanism. (Alarcón et al. 1998).

The presence of acidic proteolytic activity in the gland of Leiblein concur to what was described by Ponce et al. (1990),who showed a maximum proteolytic activity at pH 3.5, although we also observed a very slight increase of activity at pH 8. This could be related to its known function on enzyme secretions (Andrews and Thorogood 2005; Mansour-Bek 1934), but also to its important support for protein digestion through the mid-esophagus by the secretion of digestive enzymes and lubricant substances to improve digestion (Ponce et al. 1990).

### Overall evaluation of the amylase and the protease activities

The detection of high amylase activity in the digestive gland of *C. concholepas* was unexpected considering its carnivorous feeding habits. Some authors argue that amylase activity is mediated by the composition of the diet (Reimer 1982), while others suggest that the production of amylase is family-specific (Chakrabarti et al. 1995), which could be the case for isolated populations with unique feeding habits. Nevertheless, the presence of amylase in the digestive gland of loco is an indicative of its capability to degrade carbohydrates, which could be related to the degradation of structural carbohydrates and the ability to digest glycogen, an energy source commonly found in animal tissue. It is widely recognized that fishes are capable of changing their feeding habits as a strategy for food utilization and the presence of isoenzymes in these species might give them an ecological advantage (Fernández et al. 2001). A similar situation might explain the presence of amylase in mollusks but to date, changes from carnivorous to herbivorous feeding habits in molluscs have never been reported. Although, changes in dietary preferences have been observed in some introduced “loco” populations, where the main diet was replaced from barnacles (*Balanus laevis and Austromegabalanus psittacus*) to ascidians (*Pyura chilensis*) (Stotz et al. 2003)

The absence of amylase activity in the gland of Leiblein was expected since it has previously been associated to the secretion of proteolytic enzymes in muricid gastropods. It is important to notice that we mainly found the presence of acidic proteases in the gland of Leiblein, which is also concurrent with the absence of amylolytic activities. This could suggest that the food digestion starts in the esophagus and it is carried out on an acidic medium on its way to the stomach. In fact, current evidence suggests that C. concholepas retains food in the intestine for up to 16 hours, while being retained in the stomach for up to 6 hours (Stotz et al. 2003), which could indicate that the food is being under the effect of enzyme degradation during prolonged periods of time. Accordingly, these long retention times could be associated to the sub-optimal temperature conditions in which the enzymes are working in the organism (∼18 C), much lower than the optimal temperature of ∼50 C determined here, therefore, food is being exposed to digestive enzymes for longer periods to improve digestion. Proteolytic activity at low pH (3.0) was detected in both the digestive gland and the gland of Leiblein, but showing a higher specific enzymatic activity in the digestive gland compared to the gland of Leiblein, probably because it is involved in the final steps of protein degradation. Nevertheless, it must be pointed out that still significant work is made by the proteolytic enzymes secreted by the gland of Leiblein, since these enzymes execute their function from their secretion in the esophagus until they reach the stomach, where the protein degradation is complemented by the digestive enzymes located there.

The detected levels of protease activity were higher than the levels of amylase activity in the digestive gland, which could be expected in a carnivorous species and keeping in mind their digestive requirements. Similar situation has been reported in numerous carnivorous fish (Xiong et al. 2011; Lundstedt et al. 2004; Natalia et al. 2004; Sabapathy & Teo 1993). Digestion of carbohydrates is usually overlooked in carnivorous species, for not being considered as fundamental for its digestive processes. Nevertheless, amylase activities are present in the digestive system of *C. concholepas* and also in fish of different feeding habits, showing high to moderate amylase activities (Furné et al., 2005; Chan et al., 2004; Hidalgo et al., 1999), but higher in herbivorous and omnivorous. The presence of amylase in the digestive system of *C. concholepas* could certainly be associated to the need of digesting glycogen, an energy source commonly found in animal tissue. While most authors suggest that the presence or absence of specific digestive enzymes is linked to the animal feeding habits, others suggest that could be associated to phylogeny (Chan et al. 2004). Studies involving digestive enzymes in molluscs and particularly carnivorous molluscs is extremely scarce, therefore, more work is needed to identify the range of enzymes present in the digestive system of these species.

## CONCLUSION

Digestive enzymes in marine organisms have been widely studied over the years. In this study, the amylase and protease activities present in *C. concholepas* were analyzed. The major goal of this research was to understand and provide relevant information about the nutritive physiology of this interesting mollusc, but also to assess its abilities to digest alternative food sources besides its normal carnivorous diet. In view of the potential interest of cultivation and establishment of field rearing of *C. concholepas*, the shown capabilities of amylase and protease enzymes in a carnivorous muricid gastropod may be an interesting feature from the perspective of feed formulation.

## ACKNOWLEDGEMENTS

We would like to thank our professor Dr. Elena A. Uribe for her incredible support and understanding during our early years. Also we thank Dr. Victor H. Ruiz (Department of Zoology, Universidad de Concepción) for his helpful review and comments on the manuscript. No funded projects were involved in this study. All the analyses were carried out from March to September 2010.

